# Coalescent tree recording with selection for fast forward-in-time simulations

**DOI:** 10.1101/2021.12.06.470918

**Authors:** Remi Matthey-Doret

**Affiliations:** Institute of Ecology and Evolution, University of Bern, CH-3012 Bern, Switzerland

## Abstract

Forward simulations are increasingly important in evolutionary genetics to simulate selection with realistic demography, mating systems and ecology. To reach the performance needed for genome-wide simulations a number of new simulation techniques have been developed recently. Kelleher et al. (2018) introduced a technique consisting in recording the entire genetic history of the population and placing mutations on the coalescent tree. This method cannot model selection. I recently introduced a simulation technique that speed up fitness calculation by assuming that fitness effects among haplotypes are multiplicative (Matthey-Doret, 2021). More precisely, fitness measures are stored for subsets of the genome and, at time of reproduction, if no recombination happen within a given subset, then the fitness for this subset for the offspring haplotype is directly inferred from the parental haplotype. Here, I present and benchmark a hybrid of the above two techniques. The algorithm records the genetic history of a species, directly places the mutations on the tree and infers fitness of subsets of the genome from parental haplotypes. At recombinant sites, the algorithm explores the tree to reconstruct the genetic data at the recombining segment. I benchmarked this new technique implemented in SimBit and report an important improvement of performance compared to previous techniques to simulate selection. This improvement is particularly drastic at low recombination rate. Such developments of new simulation techniques are pushing the horizon of the realism with which we can simulate species molecular evolution.

## Introduction

Forward-in-time numerical simulations play an important role in evolutionary biology and ecology whether it is used to test hypothesis, to formulate expectations of the real world (Matthey-Doret and Whitlock, 2019), to investigate the consequences of potential conservation scenarios (Grummer et al., In review), as a method for inference (Excoffier et al., 2021) or to test such methods (Lotterhos and Whitlock, 2014). While coalescent based simulations have demonstrated useful in many circumstances (e.g. Excoffier et al., 2021), forward-in-time simulators offer much more flexibility and allow for modeling of selection. Study designs are often limited by the performance capabilities that simulation techniques offer. It is common for example to rescale population and genetic parameters in an attempt to speed up simulations (e.g. Booker and Keightley, 2018), however that sometimes comes at a loss of realism. Several authors have therefore worked on improving simulation techniques and hence pushing further the horizon of the kind of simulation scenarios that can be with reasonable computational resources.

In 2018, Kelleher et al. devised a simulation method for neutral loci. The method consists at recording the entire genetic tree of the population during the simulation and then, place the mutation on the tree at the end of the simulation. This simulation technique consists in storing in memory two large vectors. In the first vector, each element is an “edge”, representing a transmission of genetic material from a parental haplotype to a child haplotype. Each edge contains four values; the ID of the parental haplotype and of the child haplotype, the left-most locus transmitted and the right-most locus transmitted. The second vector matches the haplotype ID to the generation at which this haplotype existed. As lineages disappear over time, the algorithm needs to periodically “simplify” the tree in order to avoid using too much RAM. The simplification step represents an important fraction of the total run time and is described in Kelleher et al. (2018). The frequency at which the ‘simplify’ function is called also affect the trade-offs between memory usage and CPU time. A vast toolkit to track coalescent trees is implemented in the tskit software package.

The Kelleher et al. (2018) simulation technique can bring an important gain in CPU time and in memory usage compared to classical simulation techniques. This advantage is particularly important at low recombination and at high mutation rate (Haller et al. 2019 and Matthey-Doret, 2021). One obvious important limitation this simulation technique is that it can only simulate neutral loci. The Kelleher et al. (2018) simulation technique has been implemented in both SLiM (Haller et al., 209; via tskit) and SimBit (Matthey-Doret, 2021). For simple simulation scenarios at least, SimBit’s implementation appears slightly faster and uses less memory than SLiM’s implementation (Matthey-Doret, 2021). While SimBit can directly returns very helpful summary statistics, SLiM can output the tree which can later be analyzed in depth with the tskit python API.

I recently introduced a simulation technique that can bring important performance advantage but works only for loci for which the fitness effects are multiplicative among loci and among haplotypes (Matthey-Doret, 2021). Let *S*_*i*_ be the selection coefficient at the *i*^th^ locus (positive if the mutation is beneficial, negative otherwise), and let *a*_*m,i*_ and *a*_*p,i*_ be the allelic values at the maternal and paternal haplotype at the *i*^th^ locus, respectively, where the allelic values can be either 0 (wildtype) or 1 (mutated). The fitness of an individual must be defined as ∑_*i*_ (*a*_*m,i*_ +*a*_*p,i*_) (1 +*S*_*i*_). Such assumption leads to a quasi-additive relationship among alleles, esp. for alleles of small effect. For example, for a locus with beneficial mutations with *S*_*i*_*=0*.*01*, the fitnesses of the three possible are 1, 1.01 and 1.0201. I called this assumption the “multiplicative fitness assumption”.

With the multiplicative fitness assumption, a simulator can subdivide the genome into segments. The algorithm computes the fitness contribution of each segment for each haplotype. The fitness of the individual is then the product of the fitness contributions of each segment of each haplotypes. When reproducing, if a segment is transmitted without recombination, then the fitness contribution of this segment in the offspring individual is simply the fitness contribution of the parental segment multiplied by the effects of eventual new mutations. This makes fitness calculation much faster as it avoids having to look through the genetic data of each segment in order to compute fitness.

The technique has been implemented in SimBit (Matthey-Doret, 2021) with drastic advantage in terms of CPU time especially at low recombination rate. The technique does not significantly affect the memory usage though.

In this article, I am presenting a new simulation technique that can be viewed as a hybrid between the technique using the multiplicative fitness assumption already implemented in SimBit and the pedigree recording technique from Kelleher et al. (2018). The algorithm subdivides the genome in segments and uses a pedigree for each segment in order to track the genetic diversity. I will refer to it as “pedigree recording with selection”.

### Pedigree recording with selection

During a simulation, it is not essential to know the mutations each haplotype is carrying in order to model selection. Indeed, when taking advantage of the multiplicative fitness assumption, all we need to know to model selection is the fitness contribution of each haplotype. The main idea of the pedigree recording with selection technique consists at placing mutations on the pedigree to avoid all mutations in all haplotypes.

As the technique uses the “multiplicative fitness assumption”, the algorithm first needs to subdivide the genome into segments and compute the fitness contribution associated with each segment. At time of reproduction, if no recombination happens within a segment, then the fitness contribution of the segment in the child haplotype can be obtained by multiplying the fitness contribution of the segment in the parental haplotype by the be mutations. It is not necessary to know the mutations carried in a segment to infer the fitness contribution. Hence, instead of copying mutations carried by the parental haplotype, we only track the positions of new mutations, saving time for copying mutations from the parental haplotypes. When a recombination event happens in a given segment, then the algorithm needs to walk back the pedigree for both the paternal and maternal contributions in order to propagate mutations down the pedigree and reconstruct the genetic data of the recombined segment. Fitness is then re-computed by looping through the mutations carried by this newly formed segment. At the end of each generation the pedigrees for each segment is being pruned from ancestral segments that are not contributing to the current population.

#### Data structure

The data structure is fundamentally different from the one from Kelleher et al. (2018) who keeps the entire genetic history into two vectors. Here, I save the entire pedigree for each subset of the genome considered with a tree-like structure. This tree-like structure is constituted of nodes, each of which having four attributes; a pointer to the parental node, an unsigned integer of the number of children, a double-precision float number the fitness contribution and a vector of unsigned integers with the list of mutation positions.

If a node is an ancestral node or a node that came to existence with a recombination event, then the pointer to the parent is null (nullptr) and the array of mutations does not only contain new mutations but contains all the mutations that the haplotype is carrying at this particular segment.

Finally, each haplotype in the population needs a unique ID. We record a matching between haplotype ID and the array of nodes it corresponds to for both the current and the previous generation.

#### Adding a new child haplotype

A function ‘addChildHaplotype’ creates a new haplotype and take in arguments the IDs of the two parents and a vector of recombination (and segregation) points. Then, it loops through each segment and create a new node with zero children, empty of mutation and pointing to the appropriate parent node. The number of children of the parental node is incremented by one. If, at a given segment, a recombination event has happened, then the algorithm propagate mutations for both parental nodes and distribute the mutations appropriately and finally, recompute the fitness component of this node. Finally, once the node has been created, new mutations are added and the fitness component is multiplied accordingly. Pseudo-code available on Annexe 1.

#### Pruning

At every generation, the tree is pruned from its ancestors that do not contribute to modern-day population. For this, the algorithm loop through each node of each haplotype of the previous generation. If a node has zero children, then the number of children of its parental node is decremented, the node is deleted and the process is repeated for the parental node up until we find a parental node that has at least one child. Pseudo-code available on Annexe 2.

#### Propagating mutations

An important advantage of this tree-like structure presented here is for propagating mutations down a specific lineage. At every recombination event happening within a segment, the algorithm needs to propagate the mutations down the lineage. Doing so with a Kelleher et al. (2018) style of structure would require looping through the entire genetic history in order to find the ancestral nodes. With this tree-like structure however, we have direct access to the parent of each node making mutation propagation more efficient.

Method 1: In order to propagate mutations, the algorithm goes up the lineage and systematically copies the genetic content into the focal haplotype. The parent pointer of the focal haplotype is set to null as we now know its entire genetic state. As a consequence, while copying mutations down to the focal haplotype, the algorithm also deleted all nodes up until the first coalescent event (up until one node has more than one child). At the first coalescent, the number of children of the parent is decremented by one.

Method 2: In order to propagate mutations, the algorithm starts from the focal node of interest and walk up the lineage until a node has no parent anymore (until a node is either an ancestor or was born from a recombination event). As a reminder, if a node has no parent, then the attribute mutation contain all mutations, otherwise it contains only the new mutations. While walking up the lineage, the algorithm pushes pointers to children in a stack. Wigh this stack, the algorithm can now walk the lineage back down to the focal node. In doing so, the algorithm copies down the mutations to the children, decrement the number of children accordingly and remove nodes that have no more children. Pseudo-code available on Annexe 3.

### Benchmark

We benchmarked different simulation techniques with both SliM and SimBit. The techniques considered are 1) SLiM with its classic method, 2) SLiM with pedigree recording without selection (using tskit), 3) SimBit with its classic method, 4) SimBit assuming that fitness effects are multiplicative among haplotypes, 5) SimBit with pedigree recording without selection, and 6) SimBit with pedigree recording with selection (technique presented in this paper). I am only considering a single panmictic population of 5,000 diploid individuals with 10^8^ loci and a genome-wide mutation rate of 10. The genome-wide recombination rate is varying from 0 to 10 (0 to ∼1,000 cM). The recombination rate is uniform among all loci. For the technique using pedigree recording with selection, I, however, added an extra case with non-recombining chromosomes (where all recombination events are essentially just segregation). This is because the nature of the variation in recombination rate along the genome is particularly influential on the performance of this technique. For simulation scenarios that can directly incorporate selection (all except the pedigree recording without selection technique), all had a minor selection coefficient of 10^−5^.

CPU times for those benchmarks are presented in figure 1. The slowest technique is SLiM with its classic simulation technique (upper grey line) followed by the classic simulation technique of SimBit (red line). Using the multiplicative fitness assumption vastly increases performance (green line; as already seen in Matthey-Doret, 2021). The fastest techniques are the ones using the pedigree recording without selection (lower grey line for SLiM and black line for SimBit). In between classic techniques and the pedigree without selection techniques is the simulation technique using pedigree with selection presented in this paper (blue lines). The pedigree with selection technique is therefore the highest performing technique able to model selection. We observe that the detail variation in recombination rate matters with the model containing non-recombining chromosomes (lower blue line) vastly outperforming a model with uniform recombination rate (upper blue line).

**Figure 1:**
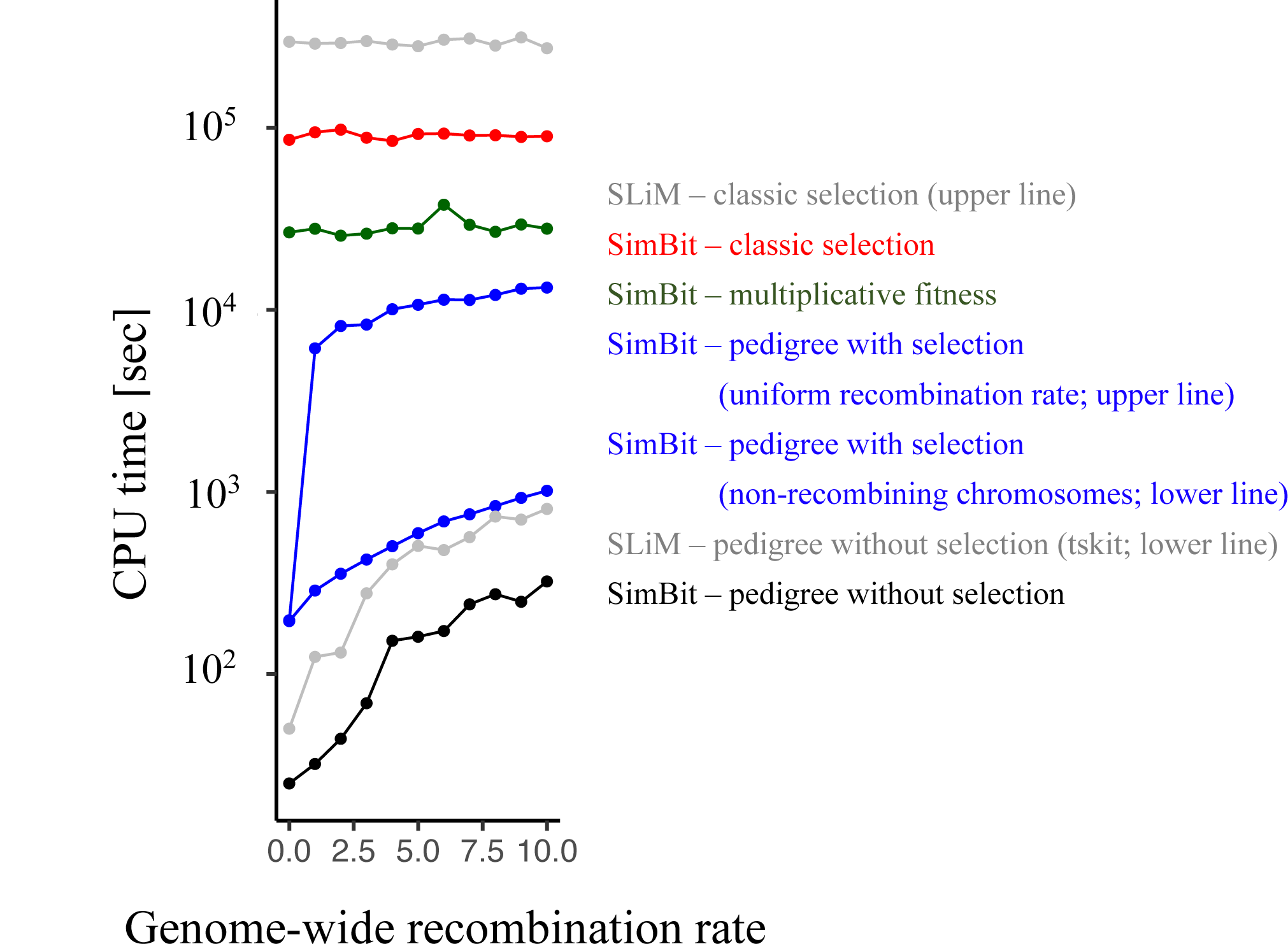
Comparison of CPU time usage for different simulation techniques with SLiM and SimBit and for varying recombination rate. In each simulation, the recombination rate is uniform among all loci, except for the lower blue line, where non-recombining chromosomes are simulated, concentrating recombination events as segregation events. The X-axis is the total number of recombination events happening, on average, over the whole genome during a reproduction event. The simulated scenario contains a single panmictic patch of 5,000 diploid individuals with 10^8^ loci and a genome-wide mutation rate of 10.

In the scenarios benchmarked, all loci were under selection. For many purposes, users want a mixture of neutral and selected loci. It is already possible in both SimBit and SLiM to track selected and neutral loci, separately. Whether this new technique will lead to higher performance in comparison to a scenario where, say neutral loci are tracked with the Kelleher et al. (2018) technique and selected ones with a classical technique (using the multiplicative fitness assumption eventually) will depend on details of the simulation parameters.

The relationship between the mutation rate and the recombination rate is an important factor influencing the relative advantage of the coalescent tree with selection technique presented here compared to a more classical simulation technique; the coalescent tree with selection performs best at low r / mu. In the benchmark, I have explored value of ratio r / mu varying from 0 to 1. For comparison, in the human genome the estimated average recombination and mutation rates are r = 7e-9 (Wang et al., 2012) and mu = 2.53e-8 (Nachman et al, 2000), hence leading to a r/mu of ∼0.28.

In this paper, I have presented a new simulation technique that can offer a drastic performance improvement compared to existing techniques. This technique is bringing us one step closer to the ability of performing realistic simulations of entire genomes.

## Supporting information

Annexe 1

Annexe 2

Annexe 3

## Data Availability

SimBit is under a permissive free program license and is available at https://github.com/RemiMattheyDoret/SimBit. Benchmark data is available in supplemental material.

