## Supplementary material for "Coalescent tree recording with selection for fast forward-in-time simulations": Annexe 1

**Annexe 1** – Pseudocode for function 'addOffspring'

This pseudocode is inspired from the function "std::pair<T8ID_type, double> T8TreeRecording::addOffspring(T8ID_type motherID, T8ID_type fatherID, std::vector<uint32_t>& recPositions);" declared and defined in files src/T8TreeRecording/T8TreeRecording.h and src/T8TreeRecording/T8TreeRecording.cpp in https://github.com/RemiMattheyDoret/SimBit.

```
Set isMother = true

Set haplotypeFitness = 1.0

for segment_index in range 0 to totalNumberSegments
(totalNumberSegments excluded)

    Set endOfSegment to the end of the 'segment_index'th segment.

    Initialize child_node = new Node

    if (recombination event happened in segment)

        propagate mutations of maternal segment

        propagate mutations of paternal segment

        Set from = beginningOfSegment # represents the first
        locus that should be copied into child_node

        while (have not finished creating child segment)

            Set from = to

            Set to = position of next recombination event or of
            end of segment

            Increase lookup to next recombination event

            if (isMother)
```

```
                    copy maternal genetic data from 'from' to 'to'
                    into child_node

               else

                    copy paternal genetic data from 'from' to 'to'
                    into child_node

               Flip value of 'isMother'.

     else

          if (isMother)

               child_node = buildChild(mother)

          else

               child_node = buildChild(father)

          if next recombination position equals end of segment

               flip 'isMother'

Mutate child_node and multiply its fitness_contribution
accordingly.

Multiply haplotypeFitness by child_node's fitness contribution

Return ID of child haplotype and haplotypeFitness
```
