## Supplementary material for "Coalescent tree recording with selection for fast forward-in-time simulations": Annexe 2

**Annexe 2** – Pseudocode for function 'prune'

This pseudocode is inspired from the function "void T8TreeRecording::prune(std::vector<std::vector<T8Segment*>>& childrenSegments);" declared and defined in files src/T8TreeRecording/T8TreeRecording.h and src/T8TreeRecording/T8TreeRecording.cpp in https://github.com/RemiMattheyDoret/SimBit.

```
For each haplotype in the population
     For each segment in the haplotype
          Set focal_node = segment
          Set parent_node = segment->parent
          While 'focal_node' has no children and 'parent_node' is
          not a nullptr
               Decrement number of children of 'parent_node'
               Delete focal_node
               Set focal_node = parent_node
```
