## Supplementary material for "Coalescent tree recording with selection for fast forward-in-time simulations": Annexe 3

**Annexe 3** – Pseudocode for function 'propagateMutations'

This pseudocode is inspired from the function "void T8TreeRecording::propagateMutations(T8Segment* focal);" declared and defined in file src/T8TreeRecording/T8TreeRecording.h and src/T8TreeRecording/T8TreeRecording.cpp, respectively in https://github.com/RemiMattheyDoret/SimBit.

```
# First, walk up the lineage and build stack of pointers toward
children in order to be able to walk down the tree

Set child_node = focal_node

Set parent_node = focal_node->parent

do

     child_node = parent_node

     parent_node = parent_node->parent

     stack.push(child_node)

while (parent has a parent)

# Walk down the lineage and copy genetic data to children,
decrement number of children and remove nodes that have only one
child.

Set parent_node = child_node

do

     child_node = stack.top() # Returns the element on top of the
stack

     stack.pop() # Remove the element on top of the stack

     child_node->parent = nullptr # Tell the child it has no parent
anymore

     Copy genetic data from parent to child

     --(parent->nbChildren) # Decrement number of children of
     'parent'
```

```
     if parent has no more children
          delete parent
     parent = child
while (child_node is not focal_node)
```
